# Noninvasive Vagus Nerve Stimulation Alters the Neural and Physiological Response to Noxious Thermal Challenge

**DOI:** 10.1101/367979

**Authors:** Imanuel Lerman, Bryan Davis, Mingxiong Huang, Charles Huang, Linda Sorkin, James Proudfoot, Edward Zhong, Donald Kimball, Ramesh Rao, Bruce Simon, Andrea Spadoni, Irina Strigo, Dewleen G Baker, Alan N Simmons

**Author notes:** **Corresponding author** Imanuel Lerman, MD, MSc, Department of Anesthesiology, University of California San Diego, 9500 Gilman Drive (0603V), La Jolla, CA 92093-0603V, USA. E-mail address.

## Abstract

The mechanisms by which noninvasive vagal nerve stimulation (nVNS) affect central and peripheral neural circuits that subserve pain and autonomic physiology are not clear, and thus remain an area of intense investigation. Effects of nVNS vs sham stimulation on subject responses to five noxious thermal stimuli (applied to left lower extremity), were measured in 30 healthy subjects (n=15 sham and n=15 nVNS), with fMRI and physiological galvanic skin response (GSR). With repeated noxious thermal stimuli a group × time analysis showed a significantly (*p* < .001) decreased response with nVNS in bilateral primary and secondary somatosensory cortices (SI and SII), left dorsoposterior insular cortex, bilateral paracentral lobule, bilateral medial dorsal thalamus, right anterior cingulate cortex, and right orbitofrontal cortex. A group × time × GSR analysis showed a significantly decreased response in nVNS group (*p* < .0005) in bilaterally in SI, lower and mid medullary brainstem, and inferior occipital cortex. Finally, nVNS treatment showed decreased activity in pronociceptive brainstem nuclei (e.g. the reticular nucleus and rostral ventromedial medulla) and key autonomic integration nuclei (e.g. the rostroventrolateral medulla, nucleus ambiguous, and dorsal motor nucleus of the vagus nerve). In aggregate, noninvasive vagal nerve stimulation reduced the physiological response to noxious thermal stimuli and impacted neural circuits important for pain processing and autonomic output.

## 1. Introduction

### 1.1 Noninvasive vagus nerve stimulation

Afferent and efferent vagus nerve signaling are critical mediators of physiological homeostasis, modulating heart rate, gastrointestinal tract motility and secretion, pancreatic endocrine and exocrine secretion, hepatic glucose production, and other skeletal and visceral functions that together make the vagus nerve the principle nerve of the parasympathetic nervous system (1). Vagal fibers can be activated with exogenous electrical stimulation carried out with surgically implanted vagus nerve stimulation (sVNS) devices (implanted around the vagus nerve in the carotid sheath). Surgically implanted vagus nerve stimulation is approved by the United States Food and Drug Administration (FDA) for the treatment of epilepsy (2) and for treatment-resistant major depression (TRMD); (3–5). However, cervical sVNS can result in complications, including hoarseness, dyspnea, nausea, and postoperative pain (6,7).

Noninvasive techniques for VNS have beneficial effects in treating epilepsy, depression, and pain. Treatment includes the use of devices that activate the auricular branch (termed Arnold’s nerve) of the vagus nerve (8–10) and the cervical vagus nerve (found within the carotid sheath) (11). Cervical transcutaneous noninvasive vagus nerve stimulation (nVNS) has shown promising therapeutic effects in the treatment of acute and chronic migraine headaches (12–14), and acute and chronic cluster headaches (15), and is now FDA-approved to treat both episodic cluster (14) and acute migraine headaches (7, 16, 17). Recent work has shown that, with finite element modeling of cervical nVNS, the electrical field significantly penetrates the human neck and is sufficient to activate the cervical vagus nerve (11). Thus, growing evidence supports the notion that transcutaneous cervical nVNS results in vagal activation that affects pain transmission and experience.

### 1.2 Pain autonomic responses and vagal nerve stimulation

Pain is a multimodal experience represented by a broad network of cortical and subcortical structures, including the primary (SI) and secondary somatosensory (SII) cortices, bilateral insular cortex (IC), anterior cingulate cortex (ACC), prefrontal cortex (PFC), thalamus, and brainstem nuclei (18, 19). Noxious thermal (painful) stimulation activates a sympathetic response, as measured by an increase in galvanic skin response (GSR); (20–22), with a dose response relationship to increasing thermal stimulus magnitude (23). Prior work has identified pain-mediated increased activation of the IC, amygdala, ACC, and PFC that correlates with pain-evoked sympathetic activity (i.e. GSR), and together offer a baseline construct for the neural basis of this autonomic pain dimension (24–28). In the present study, we used functional magnetic resonance imaging (fMRI) and primary physiological outcomes (GSR) to test the hypothesis that nVNS may alter typical cortical and subcortical neural and physiological autonomic responses to aversive noxious thermal stimuli more than to sham treatment. Prior literature supports antinociceptive effects of vagal nerve stimulation in preclinical pain models (29–34). The antinociceptive effects of VNS are postulated to depend on afferent signaling to the nucleus tractus solitarius (NTS), nucleus raphe magnus (NRM), and locus coeruleus (LC) (31). Based on this work, it has been proposed that vagal afferent inputs to NTS, NRM, and LC result in a summative signal (including activation of descending noradrenergic, serotonergic, and spinal opiodergic tracts) that inhibits dorsal horn neurons (33) (31) (34). Adding to preclinical work, multiple translational clinical studies also show similar antinociceptive effects of acute (10, 35–38) and chronic VNS (39).

Recent fMRI studies have revealed that nVNS affects brain areas important in pain processing (e.g. the medial thalamus, dorsal ACC, IC, and PFC; (40–43), thus highlighting a potential supraspinal vagal influence on pain perception. Only a single small pilot study (*n* = 20) has evaluated the neural effects of transcutaneous VNS using auricular “Arnold’s nerve” stimulation on experimental pain (36). The results did not show a difference between groups, but a post-hoc analysis of “responders”, i.e. subjects (*n* = 12) with increased pain threshold post-nVNS, showed decreased activation during the application of pain stimuli in the left dorsoposterior insula, ACC, ventromedial PFC, caudate nucleus, and hypothalamus (36). Notably, this study performed continuous transcutaneous auricular VNS during the noxious thermal challenge, possibly confounding the results as emerging literature shows pronociceptive effects during actual VNS, while the antinociceptive effects occur post-VNS (44, 45). Taken together, the evidence accumulated to date suggests that VNS alters clinical pain perception, but that VNS must be carefully timed to produce antinociceptive effects.

### 1.4 Study objectives

The objective of this study was to gain a richer understanding of post-nVNS effects on sensory discriminative neurocircuits, affective pain neurocircuits, and the peripheral autonomic response to noxious thermal stimuli. Our goal was to determine the extent of post-nVNS neural effects on pain-related brain activation and autonomic tone. Taken together, this knowledge could guide and improve the efficacious use of nVNS in pain-disease states.

## 2. MATERIALS AND METHODS

### 2.1 Participants

Thirty male and female subjects (age range, 18–54) were recruited through the Altman Clinical and Translation Research Institute at the University of California, San Diego Health System. Screening, exclusion, and inclusion criteria are found in Supplementary Information (Supplementary Information 1.1). All participants were right-handed and provided written, informed consent to participate in the study. The Institutional Review Board at the University of California, San Diego Health Systems approved this study (UCSD IRB project # 150202).

### 2.2 Intervention

Subjects were randomized to receive either nVNS (n=15) or sham (n=15) treatment (**Figure 1a**). A pair of nonferromagnetic stainless-steel surface electrodes (1-cm diameter) were placed on the subject and secured with an adjustable Velcro strap collar. The 2 devices were identical in appearance and subjects were blinded to specific intervention. Application of the device was made to either the right anterior cervical area (overlying the carotid artery) for active nVNS, or the right lateral cervical area (posterior to sternocleidomastoid) for the sham treatment. Surface electrodes were connected to the battery-powered stimulation unit by a 6-m shielded, grounded cable. Both the sham and nVNS devices delivered 1-ms duration bursts of 5 sinusoidal wave pulses at 5000 Hz with a repetition rate of 25 Hz, and a continuous train duration of 2 minutes. In both the nVNS and sham treatments, a computational fixed, initial 30-second ramp-up period was followed by 90 seconds of peak stimulation. In the nVNS treatment, the voltage was increased to 24 V, whereas in the sham stimulation it was increased to 9 V. Sham low-voltage stimulation applied to the neck far lateral to the sternocleidomastoid produces a slight tingling sensation (activating cutaneous afferents) but does not penetrate deep below the skin surface or result in muscle activation (11). Both nVNS and sham stimulation were carried out 9.5 minutes prior to the noxious thermal stimulus paradigm (**Figure 1b**).

**Figure 1.**
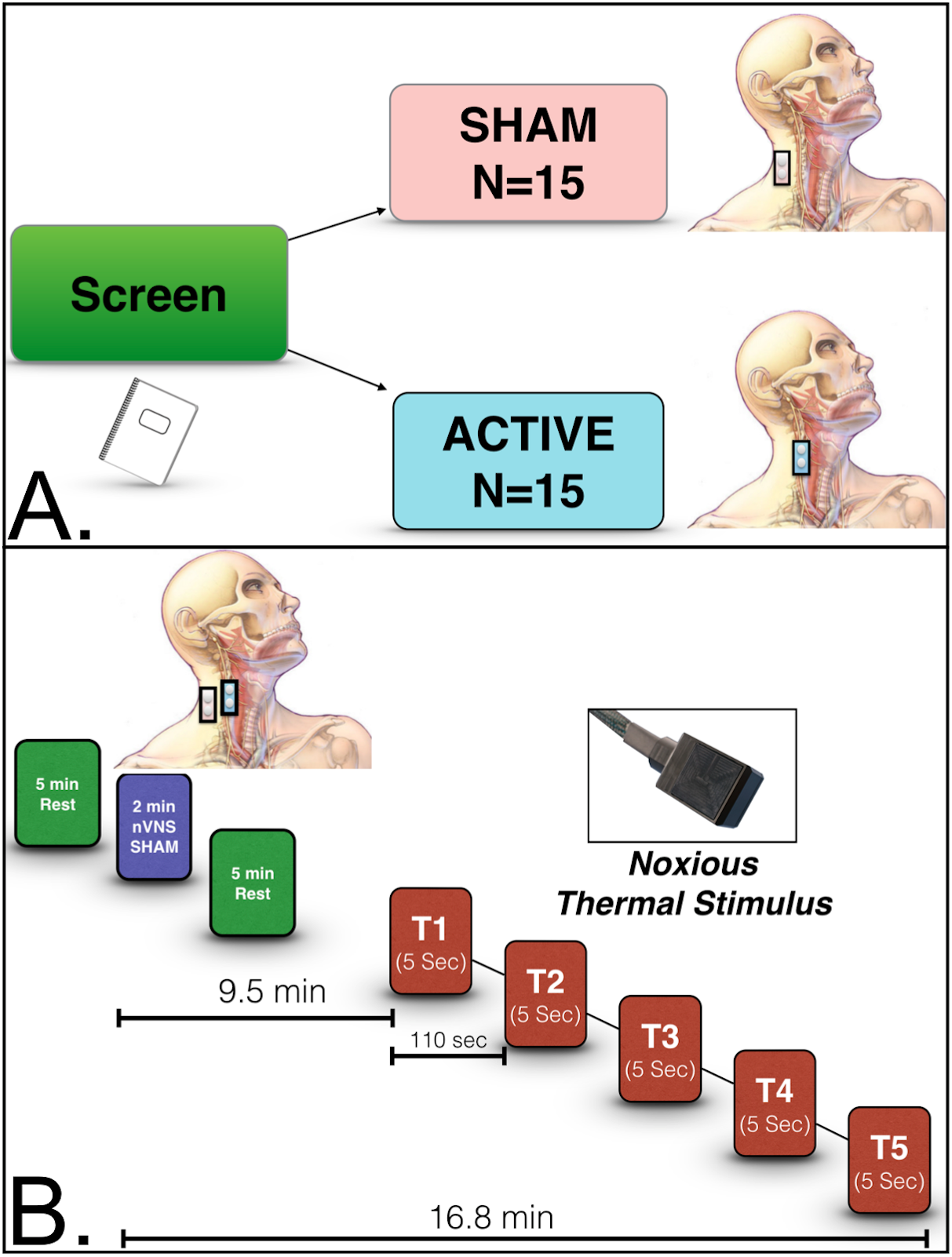
Study Design (a). Subjects were screened and randomized to either the sham treatment or nVNS group. Sham stimulation was carried out posteriolateral to the sternocleidomastoid. In the nVNS group, stimulation occurred anteromedial to the sternocleidomastoid and lateral to the trachea. In both the nVNS and sham treatments, a computational fixed, initial 30-second ramp-up period was followed by 90 seconds of peak stimulation. In the nVNS treatment, the voltage was increased to 24 V, whereas in the sham stimulation it was increased to 9 V. **Experimental design (b)**. Subjects were allowed to rest for 5 minutes before undergoing 2 minutes of nVNS (electrodes placed over carotid) or sham stimulation (electrodes placed far lateral to the sternocleidomastoid). Subjects then rested for an additional 5 minutes. Nine and a half minutes after either nVNS or sham stimulation, 5 successive noxious thermal stimuli were applied in bouts of 5 seconds each, up to 49.8°C. Each heat stimulus began 110 seconds after the start of the previous one. Measurements were taken 9.5 to 16.8 minutes after nVNS or sham.

### 2.3 Thermal stimulus task

The thermal heat threshold and thermal heat tolerance were obtained prior to the MRI scan, as previously described (46) (Supplementary Information 1.2). During the MRI scan, noxious thermal stimulation up to a temperature of 49.8°C was applied for 5 seconds via a fMRI-compatible thermode (probe size 3 × 3 cm; TSA-II, NeuroSensory Analyzer, MEDOC Advanced Medical Systems, Rimat Yishai, Israel) attached via Velcro strap, to the left lower extremity (left anteromedial lower leg, anterior to the medial gastrocnemius) in all participants. Five noxious thermal stimuli were successively applied for 5 seconds each, with a 105-second interval between each application. The total duration of the task was 9 minutes and 15 seconds (**Figure 1b**). Ten seconds after each thermal stimulus ended, each subject was asked to rate their pain intensity on the numerical pain rating scale (NPRS). In response to each thermal heat stimulus, subjects indicated the appropriate pain-intensity score with a cursor pointing to the NPRS number 0 to 10 (where 0 = no pain, and 10 = most intense pain possible). The NPRS is a validated pain-intensity score, with a test-retest reliability of 0.71 to 0.99 that is highly correlated with the numerical pain rating scale and McGill Pain Questionnaire (47).

### 2.4 Galvanic skin response

We used the BioPac MP150 Psychophysiological Monitoring System (BioPac System Inc., Santa Barbara, CA) to measure psychophysiological reactivity at rest and during the noxious thermal stimulus pain paradigm. The GSR was recorded using 2 electrodes positioned on the volar pads of the distal phalanx of the middle and ring fingers of the right hand, and was sampled with a frequency of 1000 Hz. The mean GSR (in microsiemens) prior to the application of each (#1-#5) noxious thermal heat stimulus (baseline GSR) was compared to the peak GSR response after the application of noxious thermal stimulus for each trial (#1–#5). The slope of GSR from baseline to peak was calculated (microsiemens/s). Additionally, the time (in seconds) from baseline (prior to each noxious thermal stimulus) to the peak GSR response (each post-noxious thermal stimulus) was measured and compared within and between groups. The mean GSR response was defined as the average GSR (over 25 seconds) obtained after the peak GSR was reached. Data analysis, including sample selection and artifact removal, was carried out with AcqKnowledge software (version 4.42, BioPac System Inc.) and the R statistical programming language, version 3.4.3 (48).

### 2.5 Image acquisition

T2*-weighted echo-planar images were acquired on a 3T General Electric Discovery MR 750 [Milwaukee, WI; 360 volumes, TR=1.5 s, TE=30 ms, flip angle=80°, FOV 24 cm, 64 × 64 matrix, 3.75 × 3.75-mm in-plane resolution, 30 3.0 mm (1-mm gap) ascending interleaved axial slices] using an 8-channel brain array coil. High-resolution T1-weighted FSPGR anatomical images (flip angle=8°, 256 × 256 matrix, 172 1-mm sagittal slices, TR=8.1 s, TE=3.17 ms, 1 × 1-mm in-plane resolution) were acquired to permit activation localization and spatial normalization.

### 2.6 Statistical analysis

#### 2.6.1 Group demographics of GSR analyses

Group differences in questionnaires and demographic analyses were calculated with Mann-Whitney U tests. BIOPAC system measurements of GSR were incorporated into a mixed-model regression to evaluate within-and between-group (nVNS vs sham) changes in GSR with each noxious thermal stimulus (from baseline, i.e. prior to each (#1-#5) noxious thermal stimulus to after the noxious thermal stimulus has been applied (#1-#5). The within-and between-group GSR post-thermal noxious stimulus mean value (microsiemens), time to peak (seconds), and slope from the baseline GSR to the peak (microsiemens/seconds) were compared. All statistical calculations were performed using the R statistical programming language, version 3.4.3 (48).

#### 2.6.2 MRI preprocessing

Structural and functional image processing and analysis were completed using analysis of functional neuroimages (AFNI) software (49) and R statistical packages. Echo planar images were slice-time and motion-corrected and aligned to high-resolution anatomic images in AFNI. Volumes with >20% voxels marked as outliers using 3dToutcount were censored and dropped from the analysis. For all group data points in the LME analyses 1.5 % data censor were identified as outlier. Percentage Outlier voxels in the time series were interpolated using 3dDespike. Functional data were aligned to standard space, resampled to 4-mm isotropic voxels, and smoothed with a Gaussian spatial filter (to 6 mm full width at half-maximum). Hemodynamics of the pain experience were modeled using line interpolation (3dDeconvolve/3dREMLfit modeled with TENT) for the span from the initiation of thermal heat stimulus and the following 15 seconds as modeled by 5 regressors overtime. These regressors were reconstructed to form a time series with 11 data points 1.5 seconds apart, which was used in subsequent analysis.

Group differences in the time course of Blood Oxygen Level-Dependent (BOLD) responses over the entire course of the pain experience were measured over the 5 noxious thermal applications. Time-course data were modeled using AFNI’s 3dDeconvolve TENT function. The TENT function is a linear interpolation of the hemodynamic response function over time described as piecewise linear splines. A group (nVNS or sham) × time, and (nVNS or sham) × time × GSR linear mixed-effects analysis (LME) using AFNI’s 3dLME was conducted to compare time-course data from nVNS vs sham. Effects of interest included (group × time) and (group × time × GSR) interactions, in which all were fixed effects without covariates. Group and GSR were handled as between subject factors and time was a within subject factor. Multivoxel multiple comparisons were performed by Monte Carlo simulations (using AFNI 3dClustSim modeled with 3-perameter modeling noise) to reduce the potential for false positive results. A per-voxel threshold of *p* < .001, a cluster-wise threshold of *p* < .001, and a minimum number of 14 voxels per cluster were used. The Montreal Neurological Institute (MNI) atlas was used to identify clusters. Brainstem nuclei localizations in the group × time × GSR LME were compared with graphical representations of brainstem nuclei from the Duvernoy atlas (50) and compared to prior grey and white matter brainstem maps by Beissner and colleagues (51).

## 3. Results

### 3.1 Participant demographics and psychiatric assessments

The mean age between the nVNS (24.7 ± 3.7 years) and sham group (30.7 ± 10.3 year) was not statistically different, as determined by a Mann-Whitney U test (*p* = .349). Subjects did not report having elevated anxiety, depression, or posttraumatic stress disorder (PTSD), as measured by the Beck Anxiety Index (BAI), Beck Depression Inventory 2 (BDI-2), or the PTSD Check List–Civilian version (PCL-C). Accordingly, no significant difference in mean scores between groups was noted for these measures. There were no significant differences in gender or race between the sham and nVNS groups. Two subjects failed the initial screen and were excluded from the study; one had a preexisting arrhythmia disorder (Wolf-Parkinson-White syndrome) and the other had braces (Table I). The total sample used for analysis (after exclusion of the 2 subjects who failed screening) was 15 subjects in each of the VNS and sham groups.

**Table I.** Subject demographics and psychiatric measures.

|  | **<u>Sham (*n* = 15)</u>**<br>**Mean (min, max) [%]** | **<u>nVNS (*n* = 15)</u>**<br>**Mean (min, max) [%]** | **<u>Significance</u>**<br>***p*** |
| --- | --- | --- | --- |
| Age (years) | 27.0 (18.0, 54.0) | 25.0 (18.0, 31.0) | 0.349<sup>a</sup> |
| Sex | 8M:7F [53%: 47%] | 11M:4F [73%: 27%] | 0.256<sup>b</sup> |
| Race |  |  |  |
| Asian | 5 [33%] | 7 [46%] | 0.460<sup>a</sup> |
| Black | 1 [7%] | 0 [0%] |  |
| White | 9 [60%] | 7 [46%] |  |
| Other | 0 [0%] | 1 [7%] |  |
| Excluded | 0 [0%] | 2 [14%] |  |
| BAI | 1.0 (0.0, 12.0) | 1.0 (0.0, 13.0) | 0.577<sup>a</sup> |
| BDI-2 | 1.0 (0.0, 14.0) | 2.0 (0.0, 17.0) | 0.538<sup>a</sup> |
| PCL-C | 18.3 (17.0, 28.0) | 18.3 (17.0, 28.0) | 0.469<sup>a</sup> |

BAI = Beck Anxiety Inventory; BDI-2 = Beck Depression Inventory 2; PCL-C = Posttraumatic Stress Disorder Check List–Civilian version. <sup>a</sup>=Mann Whitney U statistical test. <sup>b</sup>=Fishers exact test.

### 3.2 Pain and physiologic measures

#### 3.2.1 Baseline pain measures

Subject responses to the baseline MPQ, measured at rest prior to thermal threshold or tolerance testing, were not different between the groups (Table I). Heat thresholds, measured using the method of limits, were similar across groups (nVNS, 41.2°C ± 2.8°C; vs sham, 41.9°C ± 2.0°C; *p* = .935), as was heat tolerance, also, measured using the method of limits,(nVNS, 49.0°C ± 1.4°C; vs sham, 48.71°C ± 1.2°C; *p* = 0.467; Table II).

**Table II.** Baseline pain measures.

|  | **<u>Sham ($n = 15$)</u>**<br>**Mean (min, max)**<br>**[%]** | **<u>nVNS ($n = 15$)</u>**<br>**Mean (min, max)**<br>**[%]** | **<u>Mann-Whitney U</u>**<br>**$p$** |
| --- | --- | --- | --- |
| Adverse events<sup>a</sup> | 1 [7%] | 0 [0%] |  |
| MPQ | 0.0 (0.0, 11.0) | 5.0 (0.0, 59.0) | 0.096 |
| Heat threshold ($^{\circ}\text{C}$) | 42.2 (39.1, 46.0) | 42.4 (34.0, 48.2) | 0.935 |
| Heat tolerance ($^{\circ}\text{C}$) | 48.7 (47.1, 50.0) | 49.3 (44.7, 50.6) | 0.467 |

MPQ = McGill Pain Questionnaire.
<sup>a</sup>Unable to continue heat pain trial.

#### 3.2.2 Pain reports during the fMRI task as measured by the NPRS

During the MRI task, 5 successive noxious thermal stimuli were administered based on thermal tolerance measures, up to 49.8°C (**Figure 1b**). The pain intensity score, measured as the mean NPRS score reported during the noxious thermal stimulus paradigm, was similar between the groups for each application of thermal stimulus (**Supplementary Figure 1**). Both groups reported NPRS scores that were lower with the fifth thermal stimulus (decrease in NPRS, -0.678, ± 0.209; *t* = -3.241; *p* = .002) compared with the first stimulus. We then compared the change in mean pain report (NPRS) across each of the successive noxious thermal stimuli (T1-T5) between groups. In contrast to the nVNS group, subjects who underwent sham stimulation showed an increase in NPRS with each of the successive noxious thermal stimuli from the second to the fourth (T2-T4) (this change in pain score for each of the successive noxious thermal stimuli (T2-T4) was calculated as a slope, i.e. sham slope; 0.150 ± 0.122) vs the decrease in NPRS with each of the successive noxious thermal stimuli observed for nVNS (T2-4), (nVNS slope; -0.233 ± 0.122; p = .0301) (**Supplementary Figure 1**). One subject in the sham group was unable to complete the fifth 5-second noxious thermal stimulus due to discomfort. No other adverse events occurred during the study.

#### 3.2.3 Galvanic skin response

The GSR was recorded with each noxious thermal stimulus. The time from the onset of the each of noxious thermal stimuli to the peak GSR was measured in seconds. Mixed-model regression analyses conducted across all noxious thermal stimuli (T1–5) and between groups (nVNS vs sham) showed a significantly shorter time to peak in the nVNS group (*p* = .020; **Figure 2a**). Post-hoc comparisons between groups (with a 2-sample *t* test) revealed that subjects who underwent nVNS had a shorter time to peak GSR compared with sham subjects during the application of noxious thermal stimuli T1 and T2 (*p* < .05). Similar trends also approached significance for T3 and T4 (*p* < .09; **Figure 2a**; Supplementary Table I). We then measured the GSR slope (in microsiemens) from the baseline GSR (prior to the application of each noxious thermal stimulus) to the peak GSR (accompanying each noxious thermal stimulus) and compared how this slope changed with each of the noxious thermal stimuli (T1–5). This GSR slope decreased equally in both groups for T1 to T3 (**Figure 2b**). But in contrast to the nVNS group, which had an average decrease in slope (−0.0461 microsiemens/second) for T3 to T5, the sham group showed an increase in the average slope to peak GSR from T3 to T5 (0.049 microsiemens/seconds), with a significant between-group difference observed (group x time interaction, -0.09508; *p* = .0412; **Figure 2b**).

**Figure 2.**
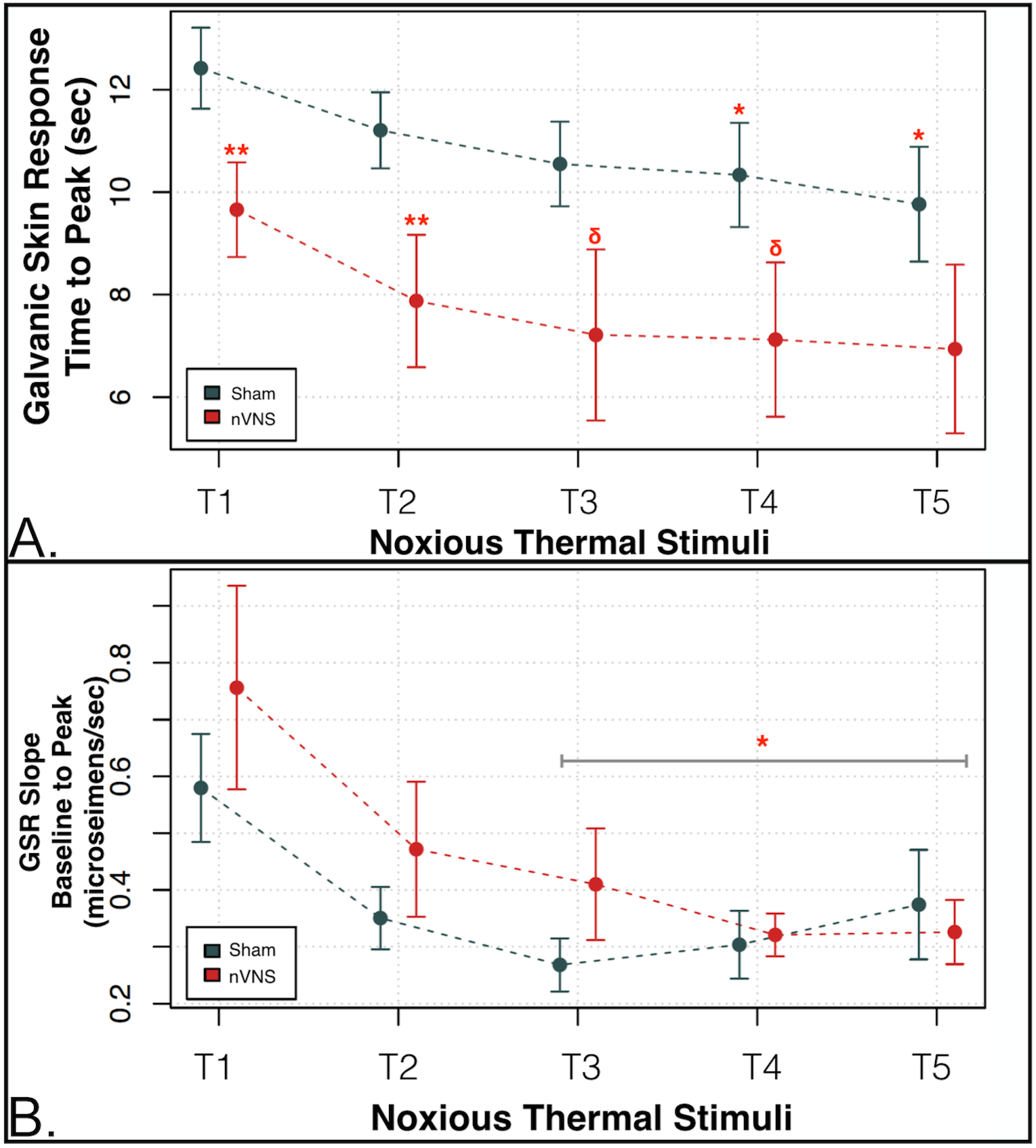
nVNS vs Sham Autonomic Measures of Sympathetic Tone Galvanic Skin Response (GSR) with noxious thermal challenge. (A) The time to peak galvanic skin response (GSR) measured in seconds after the application of each of the noxious thermal stimuli was significantly reduced in the nVNS group for noxious thermal stimuli 1 and 2 (T1 and T2) (\*\**p* < .05) compared with the sham group, and approached significance for T3 and T4 (^δ^*p* < .09). Mixed-model regression showed that the combined (T1-T5) time to peak GSR in the nVNS group was significantly shorter compared with the sham group (*p* < .02). (B) The GSR slope (in microsiemens) from the baseline GSR (prior to the application of each noxious thermal stimulus) to the peak GSR (accompanying each noxious thermal stimulus) was measured in each group. The slope from the baseline GSR to the peak response decreased in both groups with each successively applied noxious thermal stimulus from T1 to T3. However, whereas the nVNS group showed a negative average slope to peak GSR of –0.0461 from T3 to T5, the sham group showed a positive average slope to peak GSR of 0.049 from T3 to T5. The between-group difference (group x time interaction = –0.09508) for T3 to T5 was significant at \**p* < .05.

Within-group analysis conducted using a Mann-Whitney U test showed that the mean GSR (measured for each of the successive noxious thermal stimuli) was successively lower in the sham group after the application of the noxious thermal stimulus for T1, compared with T4 and T5 (*p* < .05); T2 vs T3 (*p* < .05), T4, and T5, (*p* < .001); T3 vs T4 and T5 (*p* < .001); and T4 vs T5 (*p* < .001; Supplementary Table II). In the nVNS group, the mean GSR was successively reduced after the application of the noxious thermal stimulus for T1 vs T3 (*p* = .016), T1 vs T4, and T5 (*p* < .005); T2 vs T3, T4, T5, (*p* < .001); and T3 vs T4 and T5 (*p* < .001; Supplementary Table III).

### 3.3 Imaging results

#### 3.3.1 Group differences during the application of thermal stimuli

There were no between-group differences in BOLD responses 1.5 seconds before the application of noxious thermal stimuli. During the application of a noxious thermal stimulus, 21 regions met cluster thresholds in group × time LME analyses (i.e., nVNS vs sham × time). Examination of this interaction indicated that regions in the left insula, right cerebellum/declive, and right cuneus had large clusters of greater activation (sham > nVNS). Additional regions important in the processing of thermal stimuli included the left somatosensory cortex, bilateral mediodorsal thalamus, right dorsal anterior cingulate gyrus, left supramarginal gyrus, and right medial frontal gyrus (orbitofrontal cortex [OFC]; Table III). A TENT function analysis showed significantly greater activation during the application of noxious thermal heat stimuli in the sham group in the SI (**Figure 3a, b**), SII (**Figure 3c, d, e**), left dorsoposterior insula (**Figure 3f, g**), and bilateral mediodorsal thalamus, as well as in the dorsal anterior cingulate (area 24; **Figure 3h, i, j**), and right medial frontal gyrus (OFC; **Figure 3k, l**).

**Table III.** Cluster results for group × time analysis of noxious thermal stimuli.

| Voxels | x | y | z | Within | BA | t-test | p-value |
| --- | --- | --- | --- | --- | --- | --- | --- |
| 261 | 19 | -63 | -24 | Right Cerebellum |  | 3.976308 | 0.0004 |
| 131 | -43 | -36 | 28 | Left Insula , Left Secondary Somatosensory Cortex (SII), Left Dorsoposterior Insula | 13 | 4.032758 | 0.0004 |
| 130 | 25 | -80 | 8 | Right Cuneus | 17 | 3.994926 | 0.0004 |
| 88 | 1 | -31 | 66 | Bilateral Primary Somatosensory Cortex (SI) | 3a | 3.899217 | 0.0006 |
| 56 | -18 | -72 | -24 | Left Cerebellum | 18 | 3.695176 | 0.0009 |
| 49 | 2 | -21 | -3 | Bilateral Mediodorsal Thalamus |  | 4.020902 | 0.0004 |
| 35 | 1 | -31 | 36 | Right Cingulate Gyrus | 31 | 3.785605 | 0.0007 |
| 33 | 32 | 42 | -12 | Right Orbitofrontal Cortex | 47 | 4.373772 | 0.0002 |
| 33 | -21 | -87 | 2 | Left Lingual Gyrus | 17 | 3.664152 | 0.001 |
| 26 | 7 | 8 | 44 | Right Dorsal Anterior Cingulate Gyrus | 24 | 4.024239 | 0.0004 |
| 25 | 3 | -78 | 48 | Right Precuneus | 7 | 3.816214 | 0.0007 |
| 23 | -40 | -44 | 8 | Left Superior Temporal Gyrus | 41 | 4.816714 | <0.0001 |
| 23 | -19 | -48 | 61 | Left Precuneus | 5 | 3.791053 | 0.0007 |
| 21 | -36 | -70 | -31 | Left Cerebellum |  | 4.120962 | 0.0003 |
| 21 | -21 | -38 | 12 | Left Caudate | 48 | 3.740923 | 0.0008 |
| 20 | -23 | -82 | 31 | Left Precuneus | 19 | 3.751421 | 0.0008 |
| 17 | -41 | -16 | 44 | Left Precentral Gyrus | 4 | 3.636735 | 0.001 |
| 16 | -21 | -31 | -2 | Left Parahippocampal Gyrus | 36 | 3.643388 | 0.001 |
| 16 | -18 | -1 | 15 | Left Caudate | 48 | 3.966382 | 0.0005 |
| 16 | 50 | -26 | 22 | Right Insula , Right Inferior Parietal Lobule<br>Right Secondary Somatosensory (SII) | 13 | 3.732593 | 0.0009 |
| 16 | -17 | -48 | 24 | Left Cingulate Gyrus | 31 | 4.035837 | 0.0004 |

BA= Broadmann's Area

**Figure 3.**
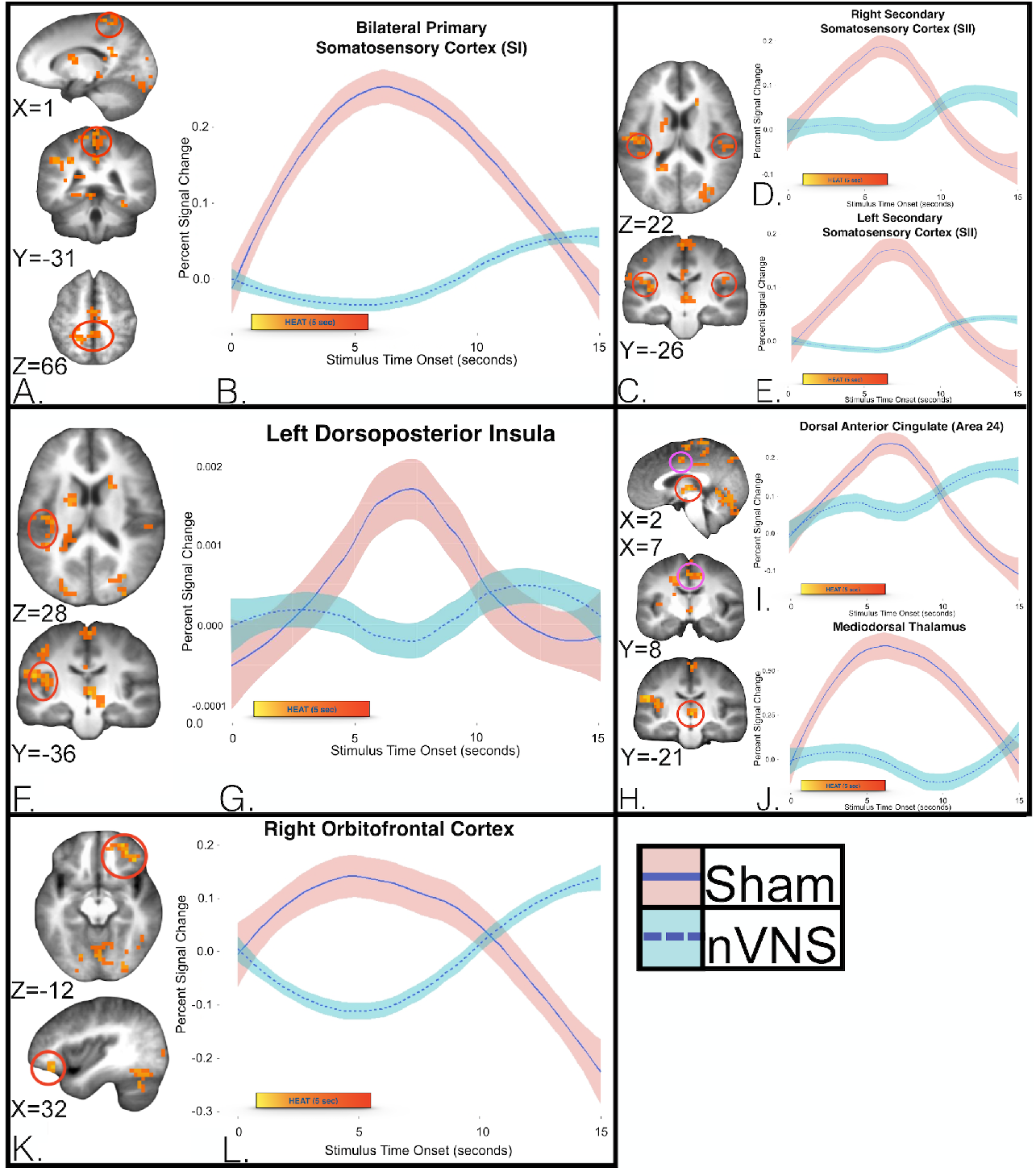
Group differences in the time course of Blood Oxygen Level-Dependent (BOLD) responses over the entire course of the pain experience. Imaging of (a) the bilateral somatosensory cortex (SI), and (c) SII, (f) left dorsoposterior insula, (h) bilateral mediodorsal thalamus and dorsal anterior cingulate (area 24), and (k) right media frontal gyrus (orbitofrontal cortex; OFC). Differential hemodynamic response curves during the application of noxious thermal stimuli 10 to 15 minutes following VNS (turquoise) and sham treatments (pink) were generated with a group × time, linear mixed-effects analysis showed that (b) subjects in the sham group had greater activity in the bilateral postcentral gyrus (SI; *p* = .0006). Treatment with nVNS significantly decreased the response of the postcentral gyrus during and after the application of noxious thermal stimuli (5 seconds each), up to 12 seconds after cessation of the painful stimulus. Subjects in the sham group had greater activity in the bilateral SII (d, e) (mean and SE shown) (right *p* = .0009, left *p =*. 0009)]. Subjects in the sham group had greater activity in the left posterior insula (g) (mean and SE shown; *p* = .0004), during and after the application of noxious thermal stimuli (5 seconds each). This result demonstrates blunting of the usual temporal dynamic response of the insula (as seen in the sham group) that is most evident during and up to 10 seconds after cessation of the painful stimulus. The sham group showed significantly greater activity in the medial thalamus and anterior cingulate (area 24) (i, j) (mean and SE shown; mediodorsal thalamus *p* = .0004, area 24 *p* = .0004), during and after the application of noxious thermal stimuli (5 seconds each). Subjects in the nVNS group had significantly decreased activity in right middle frontal gyrus (l), overlapping with the medial and lateral OFC (mean and SE shown; *p* =.0002) followed by an increase in OFC response (greater than sham) that was most evident at the 10 to 15 second mark.

#### 3.3.2 Imaging results with LME analysis

To better understand the relationships between neural and autonomic measures during thermal stimuli, the GSR mean, measured from the peak after thermal stimulus for 15 seconds, was incorporated into a group (nVNS vs sham) × linear time x GSR LME analysis using AFNI’s 3dLME to compare time-course data from the nVNS and sham groups. The group × time × GSR interaction showed that 3 regions met cluster thresholds; the postcentral gyrus/somatosensory cortex (**Figure 4a, b**), cerebellum/medullary brainstem (**Figure 4c, d**), and left occipital gyrus (Table IV). At the medullary level (i.e., level of the olive from the lower pons, spanning to the lower medulla) multiple afferent fibers enter the brainstem, including vagus, glossopharyngeal, hypoglossal, and accessory nerves that synapse on multiple brainstem nuclei (i.e., nucleus ambiguous (NAmb), dorsal motor nucleus of the vagus nerve (DMNX) and nucleus tractus solitarius (NTS)). Other brainstem nuclei important for pain processing (i.e., the rostral ventromedial medulla (RVM), rostral ventrolateral medulla (RVLM), and nucleus reticularis (Rt)) are also found at this level. Brainstem nuclei localizations were compared with graphical representations of brainstem nuclei from the Duvernoy atlas (50) and compared with prior grey and white matter brainstem maps by Biessner and colleagues (51). Subjects in the sham and nVNS groups were separated by median into high and low mean GSR categories, and the group × time × GSR interaction in the areas corresponding to the above nuclei (within medulla/brainstem) were examined. During the application of noxious thermal stimuli, subjects who underwent sham treatment and showed a high GSR demonstrated greater activity in the medulla/brainstem, compared with other groups (**Figure 4c, d**).

**Table IV.** Cluster results of group × time × GSR LME analysis.

| Voxels | x | y | z | Within | BA | *t* test | *p* value |
| --- | --- | --- | --- | --- | --- | --- | --- |
| 414 | 3 | -49 | -54 | Bilateral Medulla Cerebellum |  | 5.398 | <0.0001 |
| 25 | 4 | -44 | 63 | Bilateral Primary Somatosensory<br>Cortex (SI) | 3b | 4.200 | 0.0002 |
| 15 | -26 | -92 | -13 | Left Lingual Gyrus | 18 | 4.158 | 0.0003 |

GSR = galvanic skin response; LME = linear mixed effects.
BA= Brodmann's Area

**Figure 4.**
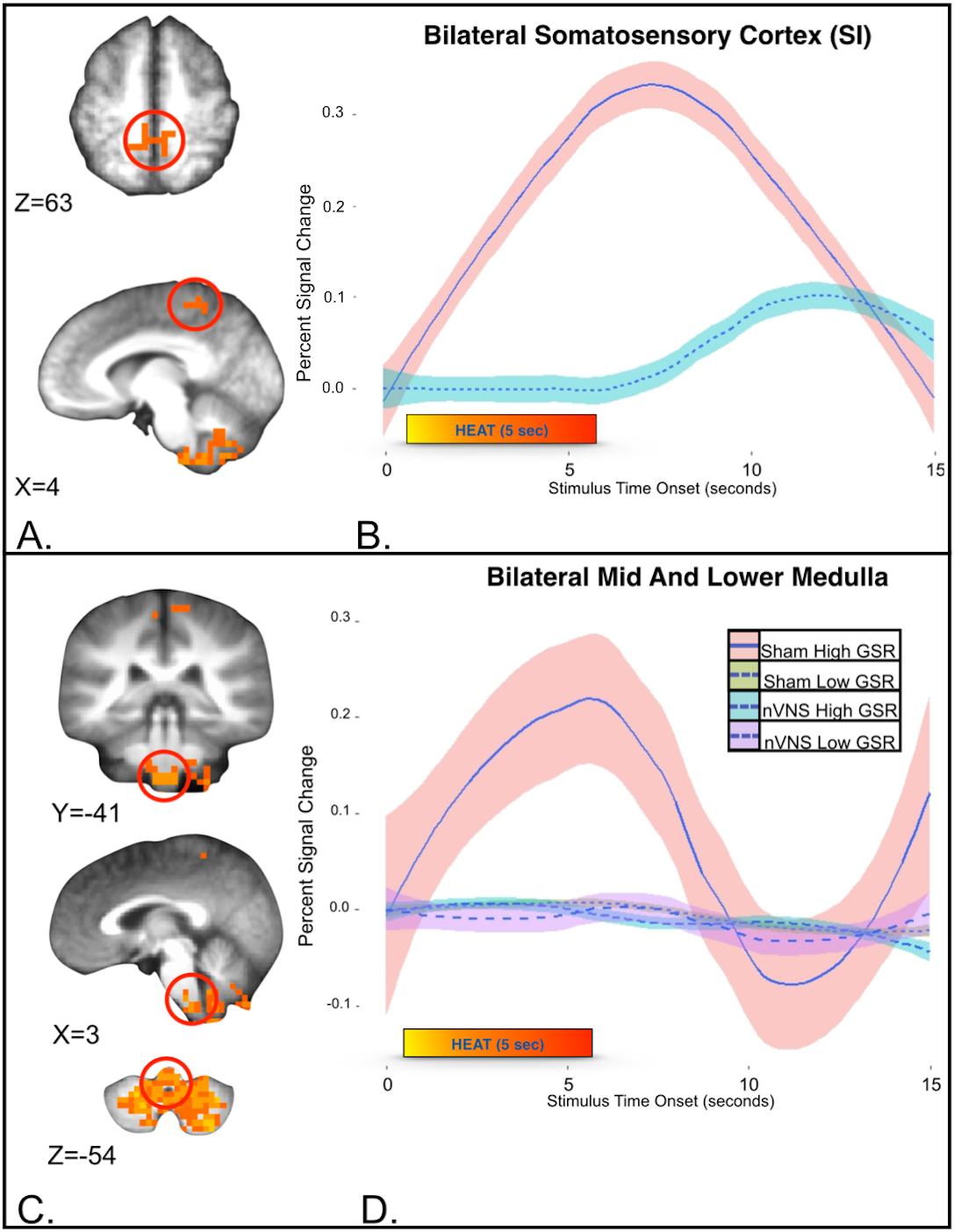
Neural and autonomic measures taken during the application of thermal stimuli (mean GSR, measured from the peak after the application of the thermal stimulus for 15 seconds). Group (nVNS vs sham) × linear time x GSR linear mixed-effects analysis. (A) Compared with subjects in the nVNS group, subjects who underwent sham treatment showed significantly greater activity in the bilateral somatosensory cortex. (B) Differential hemodynamics of pain following nVNS (turquoise) and sham (pink) treatment. (SI; mean and SE show; *p* = .0002). (C) Cerebellum/medullary brain stem measures taken during the application of thermal stimuli show (D). To assist in visual representation of this region of interest, the sham and nVNS groups were separated into high and low mean GSRs (using a group median of 16 microsiemens; the high group included 5 subjects who received sham treatment and 7 subjects who received nVNS treatment). (D) Only the high-GSR sham group (pink shade with blue line) demonstrated greater activity in the medulla/brain stem with the application of noxious thermal stimuli. At this medullary level (i.e. the level of the olive from the lower pons, spanning to the lower medulla) multiple afferent fibers enter the brainstem, including the vagus, glossopharyngeal, hypoglossal, and accessory nerves, that synapse on multiple brainstem nuclei [i.e. the nucleus tractus solitarius (NTS), nucleus ambiguous (NAmb), and dorsal motor nucleus of the vagus nerve (DMNX)]. Other brainstem nuclei important for nociception [i.e. the rostral ventrolateral medulla (RVLM), rostral ventromedial medulla (RVM), and nucleus reticularis (Rt)] are also found at this level.

## 4. Discussion

The effects of VNS on the central and peripheral neural circuits involved in pain and autonomic physiology are not well elucidated. In this study nVNS treatment (when compared to sham) resulted in reduced responses in highly relevant pain-processing nodes. There was a significant alteration of autonomic tone, as determined by a decrease in sympathetic activity (measured with GSR) and attenuated activity in brainstem nuclei known to contribute to pain-mediated autonomic responses. These results provide preliminary evidence of significant nVNS modulation of central and peripheral autonomic neural circuits relevant to pain perception.

### 4.1.1 Post-Non-invasive Vagus Nerve Stimulation With Noxious Thermal Stimuli; Neural Effects on Bilateral Somatosensory Cortex 1 (SI) and Somatosensory Cortex 2(SII) (i.e., Lateral Pain Pathway)

Compared with subjects in the sham group,group × time LME analysis showed that subjects in the nVNS group had decreased neural activation of SI and SII, the medial dorsal thalamus, ACC, IC, and OFC—all brain regions associated with the processing of painful stimuli. Meta-analysis of human data from fMRI, EEG, magnetoencephalography (MEG), and positron emission tomography (PET) studies has shown that the commonest regions found to be active during an acute pain experience (18) are the SI and SII, thalamus, ACC, IC, and PFC, (comparable to areas that show decreased activity with nVNS in this study). Analysis of the group × time interaction showed a decrease in responses of the bilateral SI and SII somatosensory cortex, suggesting that nVNS mediates this signaling during the application of a thermal stimulus. These nVNS-mediated response changes in the SI somatosensory cortex strip match bilateral somatotopy to the lower leg, consistent with the placement of the Peltier heat probe. It is generally believed that somatosensory stimuli are processed primarily or preferentially by the hemisphere that is contralateral to the point of stimulation. However, evidence from clinical studies in patients with brain lesions, and from brain-imaging studies of noxious painful stimuli have called this theory into question (52). Well-established brain regions that show bilateral activation upon the application of painful stimuli include the ACC, PFC, SII, insula, thalamus, and inferior parietal lobe (53–58); and, in some instances, SI (53, 59–61). It is likely that nVNS-mediated bilateral decreases in SI represent modulation of cortical context and or anticipatory neurocircuits. We postulate that the observed effects of nVNS on bilateral pain-processing pathways may represent bilateral nVNS afferent signaling effects; possible afferent to bilateral efferent effects on the thermal (and possibly nociceptive) signaling pathways of the spinal cord; or direct disruption of normal bilateral thermal and nociceptive afferent neural firing patterns that either independently or collectively change the temporal dynamics of pain processing.

### 4.1.2 Post-Non-invasive Vagus Nerve Stimulation With Thermal Stimuli; Neural Effects on Left Dorsoposterior Insula

In addition to nVNS-mediated bilateral SI and SII responses, our analysis showed a unilateral decrease in the left dorsoposterior insula. The dorsoposterior insula exhibits an anterior-to-posterior somatotopic organization in response to innocuous or noxious/painful stimuli as measured with fMRI (62–66). Various painful stimuli, including hypertonic saline injection (63), thermal stimuli (64), and laser stimuli (67), have consistently reproduced this anteroposterior somatotopy within the dorsoposterior insula; specifically, rostral targets (head/neck) localizing more anteriorly whereas caudal targets (leg) localizing posteriorly (68). Dorsoposterior insular stroke results in discrete thermoanesthesia and analgesia that equivalently mapped anteroposterior somatotopy, further supporting the idea that the dorsoposterior IC plays a critical role in the pain experience (69–73). Neuroanatomical data have demonstrated that the lamina I spino-thalamo-cortical pathway convey both nociceptive and interoceptive information mapped to the viscerosensory cortex in the posterior and mid-insular cortex, which is then represented in the anterior insula (74–76). Surgically implanted vagal nerve stimulators (FDA-approved for treatment of resistant depression and epilepsy) consistently (77–82) modulate insular cortex activity, thus pointing to the insula as a possible neuromodulatory target for nVNS. Moreover, while insular activity is known to increase during acute VNS (77–79), recent work has shown a resultant decrease in insular activity at 10 to 15 minutes post-nVNS (42). In our cohort, there was a significant left dorsoposterior insula decrease in activity 10 to 17 minutes post-VNS that further support the temporally dependent dose-response effects of VNS.

As a whole, the observed changes in the response to pain in the SI, SII, and left dorsoposterior insula with nVNS infer possible nVNS-mediated changes in neuronal firing patterns, either through direct brainstem effects, afferent cortical, or afferent cortical-to-efferent brainstem/spinal cord effects on nociceptive signaling.

### 4.1.3 Post-Non-invasive Vagus Nerve Stimulation With Thermal Stimuli; Neural Effects on Bilateral Mediodorsal Thalamus and Anterior Cingulate Cortex (Area 24) (i.e. Medial Pain Pathway)

Beside lateral thalamic nuclei projections (i.e., ventroposterior-lateral and ventroposterior-medial thalamic nuclei) to the SI and SII, known to relate the sensory-discriminative aspects of pain spinal pathways to limbic structures, the medial thalamic nuclei provide inputs to emotion-related brain areas, including the insula, ACC, amygdala, PFC, and other regions important in processing the affective-motivational dimension of the unpleasant pain experience (83). In our study, the nVNS group showed a decreased response in the bilateral mediodorsal thalamus and dorsal ACC (Brodmann area 24) during the application of thermal stimulation, (with the group × time interaction). Prior clinical work shows that the mediodorsal thalamus is important in antinociceptive regulation (84), the processing of emotions (85), affective pain processing (pain unpleasantness) (64, 86, 87) (88–90),(85), thought to occur through mediodorsal thalamic connections with dorsal ACC (area 24). In an illustrative case study, a patient with a somatosensory cortex stroke that spared the dorsal ACC (area 24) and thalamus (including mediodorsal thalamus) reported usual contralateral limb analgesia to painful stimuli, but the patient continued to reported an “unpleasant” feeling with the application of painful stimulus, suggesting *in vivo* separation of the affective and sensory discriminative pain pathways (87). We observed mediodorsal thalamus and dorsal anterior cingulate deactivation in the nVNS group, which likely indicates a key mechanism of the effect of nVNS on the medial affective pain pathway, in agreement previous studies (84, 91, 92). Based on this remarkable (but preliminary) finding in a future study we will measure nVNS effects on affective pain (i.e. pain unpleasantness).

### 4.1.4 Post-Non-invasive Vagus Nerve Stimulation With Thermal Stimuli; Neural Effects on Right Orbitofrontal Cortex

In addition to the medial dorsal thalamic connections to ACC, there are known medial thalamic projections to the PFC, ventromedial-prefrontal, and orbitofrontal (OFC) cortices (93, 94). The group × time analysis in the current study showed decreases in the right OFC response, suggesting that nVNS mediates this signaling during nociceptive stimulation (**Figure 3 k, l**). Prior clinical work has also demonstrated involvement of the prefrontal and frontal cortical regions in reflecting the emotional, cognitive, and interoceptive components of pain conditions, negative emotions, response conflicts, decision-making, and appraisal of unfavorable personal outcomes (95, 96). Multiple pain-imaging studies have found that the frontal cortical regions are critical for controlling functional interactions among key brain loci that produce changes in the perceptual correlates of pain, independent of changes in nociceptive inputs (64, 97, 98). Manipulating the cognitive aspects of pain, such as reappraisal, control, and coping, produce neural changes in the brain thought to be important in top-down processing. The lateral OFC expresses a contextual modulation of response that is widely implicated in emotional regulation and decision-making behaviors (99, 100), and it has been postulated that the valuation of pain is context-sensitive, as classified by the OFC (101). Activity in the ventromedial cortex and the OFC has repeatedly been shown to be modulated by acute (78, 102-105) and chronic VNS (78, 102, 106). In this cohort, we showed initial decrease in OFC activation during nociceptive thermal stimulation followed by an increase in OFC response (greater than sham) that was most evident at the post-thermal stimulation 10 to 15 second mark. This interesting finding suggests a decrease in the OFC affective appraisal of pain (0-6 seconds) followed by a subsequent late hemodynamic response increase that may reflect a resultant increase in pain-coping behavior. The observed nVNS-mediated decrease in the response of the OFC during the application of maximal noxious thermal stimulation is consistent with the results of prior VNS-treatment imaging studies, and suggests that the effect of nVNS on the OFC likely plays a role in the processing of painful and aversive stimuli.

### 4.1.5 Non-invasive Vagus Nerve Stimulation; Combined Neural Effects and Physiological Measures (GSR)

The group × time × GSR analysis highlighted differential interactions among nVNS, GSR, and the temporal dynamic of pain responses in the cerebellum, medulla/brainstem nuclei, bilateral SI, and a right occipital gyrus cluster. In addition to cortical nodes, the mid and lower medullary brainstem have been shown to be important sites that demonstrate an interaction between sympathetic output and pain, with decreases in sympathetic output (as measured with cardiac vagal tone) shown to correlate with brain stem nuclei including: 1) RVLM, 2) Rt, 3) NAmb, 4) DMNX and 5) the RVM, (all found superior to the obex at the level of the olive spanning to lower medulla) (107). In this study, medulla/brainstem clusters from sham and nVNS groups were separated into high and low mean GSRs. Only the sham treatment group showed a high GSR, demonstrated by greater activity in the medulla/brainstem, compared with other groups (sham low, nVNS high, nVNS low). At the level of the medulla, where this interaction is found (i.e., superior to the obex at the level of the olive from the lower pons, spanning to the lower medulla) multiple afferent fibers enter the brainstem, including the vagus nerve, and the glossopharyngeal, hypoglossal, and accessory nerves, as well as multiple nuclei and tracts (i.e., DMNX, NTS, NAmb, RVM, RVLM, and the Rt). In particular, the Rt is proposed to be primarily a pronociceptive center that integrates multiple excitatory and inhibitory functions important in nociceptive processing (108). The premotor nuclei (i.e., NAmB and DMNX) are critical in autonomic response patterns evoked by physiological and sensory stimuli (109) that culminate in efferent parasympathetic outflow and play a crucial role in parasympathetic reflexes, accepting input from the NTS that is the principal nucleus for incoming afferent signals from the vagus nerve (110). The RVM is intricately involved in areas of endogenous pain modulation in the brain, conveying descending pain modulatory influences from the PAG to neurons located in the dorsal horn of the spinal cord. The ON and OFF cells of the RVM increase or decrease activity during the application of painful stimuli, respectively (111), with notable effects on descending pain-inhibitory circuits (112). In sum, decreased activity in the medulla found in this study can be seen in reduced autonomic tone (reduced GSR in the lower GSR sham group), or through vagal nerve stimulation (in both nVNS groups, regardless of GSR (high or low)) suggesting that the default regulation of GSR can be decoupled through nVNS. We postulate that, even with increased GSR output (high GSR in the nVNS group), nVNS inhibits the response of the central nervous system to pain (in part) by blunting the response in key nuclei in the medulla that relay autonomic responses. Support for the relationship between the nVNS neural response and physiological response stems from altered autonomic sympathetic output (i.e., time to peak GSR and decrease in GSR slope). Future study is planned to examine this interaction in disease states such as Posttraumatic Stress Disorder or Major Depressive Disorder where dysfunctional emotional regulation and dysregulated autonomic output coincide.

### 4.2.1 Non-invasive Vagus Nerve Stimulation; Autonomic Measures and pain report

Time to peak GSR (i.e., time from GSR measured immediately prior to each 5 second noxious thermal stimuli to peak post-noxious thermal stimuli) in the nVNS group was more rapid than in the sham group, indicating changes in the temporal dynamics of pain processing and subsequent sympathetic output. The temporal dynamic of GSR during the application of a thermal stimulus is an important component of autonomic responsivity (20–22), and the subsequent emotional regulation of aversive stimuli (113, 114). Loggia and colleagues demonstrated the existence of a dose-response relationship between the magnitude of a thermal stimulus and the time to peak GSR (23). Specifically, their study showed that the greater the impact of a stressor (increased thermal temperature), the greater the rise in GSR, thus resulting in a longer time to peak response. In addition to the longer time to peak observed in the sham group, significant differences between the nVNS and sham groups in the slope of the GSR rise from baseline (prior to each noxious thermal stimuli to peak after noxious stimuli) for the latter thermal stimuli (T3-T5) were observed. In particular, the slope of the GSR response decreased across the length of the task in the nVNS group, whereas the slope of the response in the sham group increased. Taken together the longer time to peak and increase in GSR slope in the sham group compared to the nVNS group further suggest nVNS alters sympathetic output, possibly to due to the brainstem and cortical effects described.

Together with group differences in GSR (time to peak and slope) there was also a significant difference in the change in response between subjects who underwent nVNS vs sham stimulation across thermal stimuli (T2-T4) in reports of pain, as measured by the NPRS. The group that underwent sham stimulation showed a progressive increase in NPRS (across thermal stimuli T2–4), whereas the nVNS group demonstrated a significant decrease in NPRS (across thermal stimuli T2–4). To further characterize the effects of nVNS, additional work is needed that carefully measures affective pain, such as unpleasantness and catastrophizing, associated with the application of noxious thermal stimuli.

### 4.4 Non-Invasive Vagus Nerve Stimulation Potential Temporal Dependent Effects on Brain & Pain

Henry and colleagues (77) first argued (2002), that neural effects which occur during VNS are very different from those that occur after VNS, while others continue to confirm this phenomenon (42, 43, 115). In this study, we showed that subjects in the nVNS group had nVNS-mediated activity decreases in the dorsoposterior insula, low medullary brainstem, medial thalamic, and ACC compared with subjects in the sham group (occurring 10 to 17 minutes after nVNS treatment). Similar to our post-nVNS effects observed on the low medullary brainstem, Frangos and colleagues also show post-cervical transcutaneous VNS effects in this time frame, (13–15 minutes after stimulation), in posterior insula, lower medullary brainstem and medial thalamic/ACC deactivation at rest (43), that provide a convergence of preliminary evidence supporting a temporal nVNS dose-response curve (43). In line with the aforementioned post-VNS neural effects, emerging clinical literature also demonstrate post-VNS antinociceptive effects (39, 116) while pronociceptive effects during VNS have also been reported(44, 45). Both prior literature and this study suggest that the temporally dependent neural effects (i.e. during vs post-stimulation) of VNS may be critical to clinically relevant pro-or anti-nociceptive effects of VNS treatment, and therefore should be taken into account in future clinical study designs. Moreover, future studies are planned to determine the temporal dose response curve on affective pain processing that may also be of clinical import to guide efficacious use of VNS for clinical comorbid pain and psychiatric syndromes.

## 5. Limitations

Our work has some important limitations. The study was carried out in healthy control subjects. Because, as a pilot study, we involved only 15 subjects per group, the small sample size may not adequately represent a larger population. Therefore, our results as described here should be considered preliminary. However, the positive findings observed in this small cohort of healthy control subjects were robust and significant, warranting further investigation of the effects of cervical transcutaneous nVNS on the brain in a larger cohort of healthy control subjects, and in subjects who may experience a greater magnitude of affective pain subtypes, that may include Posttraumatic Stress Disorder or Major Depressive Disorder. Our study found significant neural alterations in the temporal dynamics of noxious thermal-stimuli processing known to be important in affective pain processing, and group differences in changes in the subjective pain report across the thermal stimuli (T2-T4). But we did not detect a difference in subjective reports of pain for each thermal stimulus with a near maximal noxious thermal stimulus (**Supplementary Figure 1**). We chose a near maximal noxious thermal stimulus to ensure clear autonomic responses (GSR). Our own work (46) as well as that of other studies has described maximal noxious stimuli that result in maximal reports of pain and, therefore, blunting of group differences in mean reports of pain (i.e. a ceiling effect on pain report) (117–119) (120). This phenomenon also could have occurred in this study. Future studies that measure affective pain (such as pain unpleasantness and catastrophizing) using maximal and submaximal noxious thermal stimuli are now needed to further characterize the antinociceptive effects of nVNS, as measured by reports of pain. While we correct for motion artifact at the brainstem level with the (Group x time x GSR) interaction, this area can be artifact-prone due to motion. Although others have shown a similar pain and autonomic tone interaction at the same medullary brainstem level (Sclocco and colleagues (107)) future study is planned in larger cohorts to confirm this interaction at this brainstem level.

## 6. Conclusion

We examined the neural effects of nVNS during a noxious thermal stimulus challenge, in the context of autonomic responses. We demonstrated 3 major findings; first, nVNS activity not only reduces peak responses to thermal stimuli in the SI, SII, medial thalamus, dorsal anterior cingulate (area 24), dorsoposterior insula, and OFC, which are important nodes in sensory discriminative pain, affective emotional pain, and interoception pathways, but also changes temporal dynamic responses within these nodes. Second, nVNS alters autonomic responses to noxious thermal stimuli, as measured by GSR, and therefore affects critical autonomic pain networks. Third, even with a higher GSR response being provoked by the application of noxious thermal stimulus, nVNS decreased the central nervous system response by blunting the usual reactions in key nuclei in the medulla that relay autonomic responses. These significant findings may improve effectual nVNS that, if tuned with careful dose-response curves in mind, could translate into efficacious targeted effects on pain and autonomic neural circuits.

## Acknowledgments

This study was funded by the VA San Diego through the Center for Stress and Mental Health. All listed authors meet the criteria for authorship set forth by the International Committee of Medical Editors. Imanuel Lerman MD MSc was the principal investigator; participated in the design of the study, recruitment and follow-up of subjects, critical review and discussion of the final study report, data collection and interpretation, and drafting of the study report and manuscript; and had final responsibility for the decision to submit for publication. All authors were involved in the interpretation, drafting, and review of the manuscript. James Proudfoot served as the study statistician. All authors provided input to the report and approved the final version.

## Supplementary material

### 1.1 Inclusion and exclusion criteria

All subjects were instructed to refrain from taking any over-the-counter analgesics or anti-inflammatory medications, tobacco products, or alcoholic beverages for 1 week prior to the study visit. All subjects underwent phone screening up to 1 week prior to the fMRI scan. Subjects with prior surgery or abnormal anatomy at the treatment site (ie, anterior cervical neck region); injury; or abnormal anatomy at the MEDOC Peltier probe site [fMRI-compatible thermode, probe size 3 × 3 cm; TSA-II NeuroSensory Analyzer, MEDOC Advanced Medical Systems, Rimat Yishai, Israel), the left, lower leg anterior shin area; a history of neurologic disease (including transient ischemic attack, seizures, and syncope); a history of any type of implanted neurostimulator device, or cardiac pacemaker; or a history of cardiovascular disease or carotid artery disease were excluded from the study. Participants had no history of eating disorders and no current Diagnostic and Statistical Manual of Mental Disorders, 4^th^ edition (DSM-IV) Axis I psychiatric illnesses per self-report, or on standardized measures.

### 1.2 Heat tolerance and threshold measurements

Thermal thresholds and heat tolerance were calculated using the previously described method of limits (Yarnitsky & Sprecher, 1994) by taking the average of 5 thermal stimuli successively applied with a fMRI-compatible MEDOC probe to the right lower extremity (anterior shin) at an increasing slope of 1°C/s, from 32°C up to a maximum of 50°C.

### 1.3 Correlations of interest

#### 1.3 Correlations between autonomic tone and pain reports

Autonomic measures taken during the application of thermal stimuli (time to peak GSR, mean GSR) and pain score were correlated within each group to better understand the relationships among these factors. In the nVNS group, mixed-model regression for the change in mean GSR with covariates for thermal stimuli showed significant negative correlations between mean GSR and thermal stimuli T3 (−0.955 ± 0.271; *t* = -3.528; *p* < .001), T4 (−1.429 ± 0.85; *t* = -5.106; *p* < .001), and T5 (−1.593 ± 0.295; *t* = -5.394; *p* < .001), that support the difference from baseline GSR to the peak slope (ie, a decrease in slope from T3-T5 in the nVNS group only). Lastly, mixed-model regression results (for the nVNS group only) showed that the time to peak GSR covaried with thermal stimulus trial and pain score, with a significant effect of pain score detected (1.035 ± 0.449; *t* = 2.305; *p* = 0.025).

**Supplementary Figure 1.**
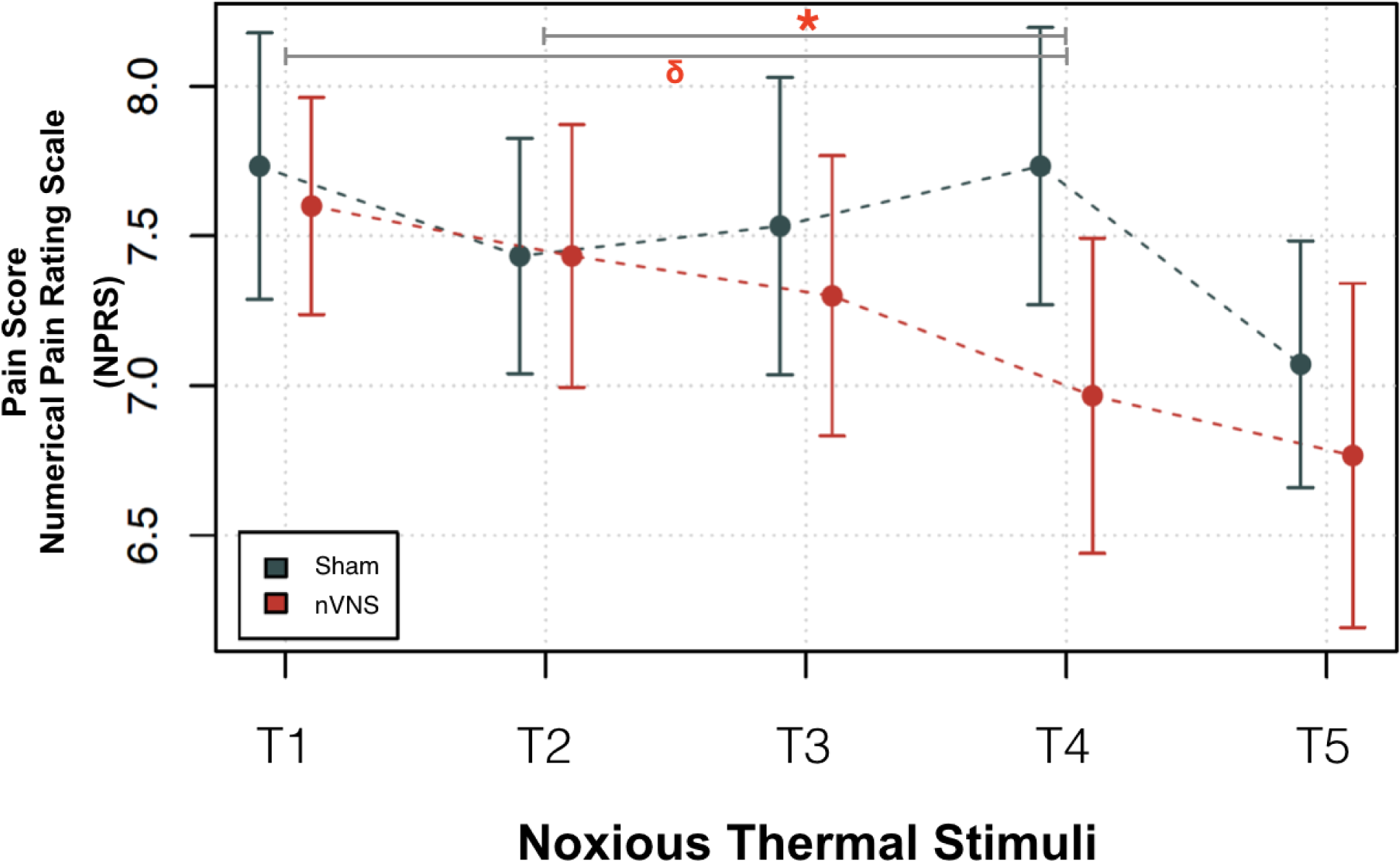
After either nVNS or sham stimulation, 5 successive noxious thermal stimuli were applied (up to 49.8°C) for 5 seconds each (T1-T5). Mean pain, as reported by subjects using the numerical pain rating scale (NPRS) after each noxious thermal stimulus did not differ between the sham and nVNS groups. Both groups had lower NPRS scores at T5 compared with T1 (NPRS decreased by –0.678 ± 0.209; *t*= –3.241; *p* = .002). In contrast to findings for the nVNS group, subjects who underwent sham stimulation had a positive slope in NPRS scores across thermal stimuli (i.e. the change in NPRS score with successive noxious thermal stimuli T1-T5) for T2 to T4 that was significantly different (slope in the sham group, 0.150 ± 0.122; vs the slope in the nVNS group, –0.233 ± 0.122; *p* = .0301) and also approached significance from T1 to T4 (sham group, 0.010 ± 0.847; vs nVNS group, –0.203 ± 0.847; *p* =.0785). Red circles = nVNS group. Blue circles = sham group. \**p* < .05;^δ^*p* < .08.

**Supplementary Table I:**
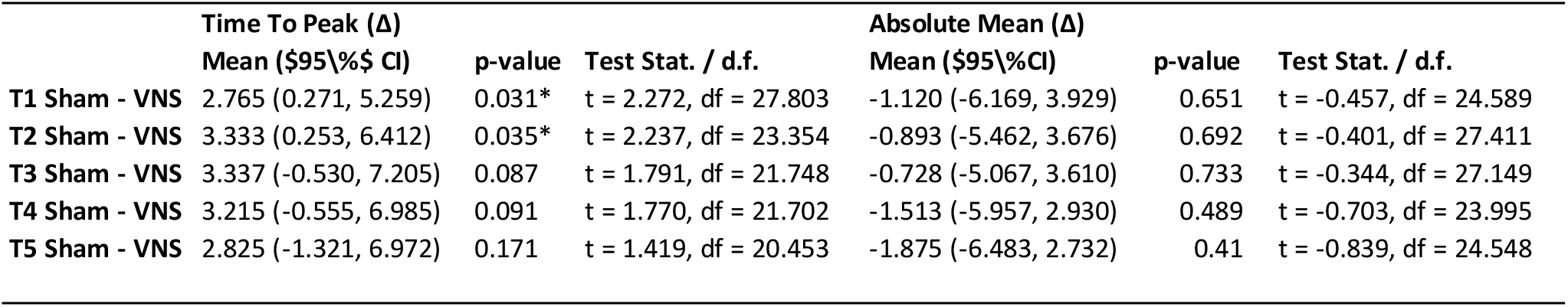
Between-group comparisons for the time to peak and absolute mean GSR. The nVNS group showed significant decreases in the time to peak GSR for T1 and T2.

| | Time To Peak ($\Delta$) | | | Absolute Mean ($\Delta$) | | |
| --- | --- | --- | --- | --- | --- | --- |
| | Mean (\$95\%\$ CI) | p-value | Test Stat. / d.f. | Mean (\$95\%\$ CI) | p-value | Test Stat. / d.f. |
| T1 Sham - VNS | 2.765 (0.271, 5.259) | 0.031* | t = 2.272, df = 27.803 | -1.120 (-6.169, 3.929) | 0.651 | t = -0.457, df = 24.589 |
| T2 Sham - VNS | 3.333 (0.253, 6.412) | 0.035* | t = 2.237, df = 23.354 | -0.893 (-5.462, 3.676) | 0.692 | t = -0.401, df = 27.411 |
| T3 Sham - VNS | 3.337 (-0.530, 7.205) | 0.087 | t = 1.791, df = 21.748 | -0.728 (-5.067, 3.610) | 0.733 | t = -0.344, df = 27.149 |
| T4 Sham - VNS | 3.215 (-0.555, 6.985) | 0.091 | t = 1.770, df = 21.702 | -1.513 (-5.957, 2.930) | 0.489 | t = -0.703, df = 23.995 |
| T5 Sham - VNS | 2.825 (-1.321, 6.972) | 0.171 | t = 1.419, df = 20.453 | -1.875 (-6.483, 2.732) | 0.41 | t = -0.839, df = 24.548 |

**Supplementary Table II:**
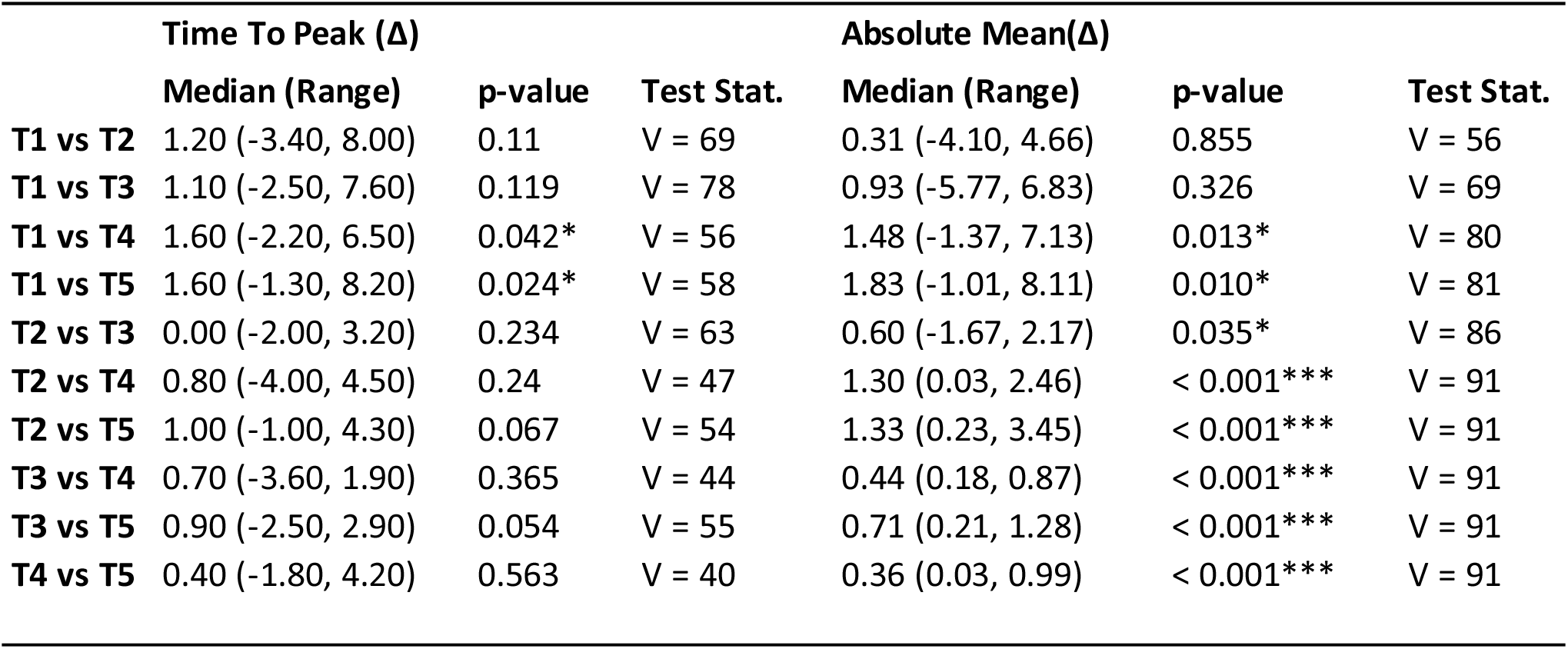
Within-group comparisons for the time to peak GSR and absolute mean GSR for the sham stimulation group. In the sham group, the time to peak GSR increased from T1 to T4 and T5. The mean GSR measured after the application of noxious thermal stimuli consistently increased from T2 to T5 and from T1 to T4.

| | Time To Peak ($\Delta$) | | | Absolute Mean($\Delta$) | | |
| --- | --- | --- | --- | --- | --- | --- |
|  | Median (Range) | p-value | Test Stat. | Median (Range) | p-value | Test Stat. |
| T1 vs T2 | 1.20 (-3.40, 8.00) | 0.11 | V = 69 | 0.31 (-4.10, 4.66) | 0.855 | V = 56 |
| T1 vs T3 | 1.10 (-2.50, 7.60) | 0.119 | V = 78 | 0.93 (-5.77, 6.83) | 0.326 | V = 69 |
| T1 vs T4 | 1.60 (-2.20, 6.50) | 0.042* | V = 56 | 1.48 (-1.37, 7.13) | 0.013* | V = 80 |
| T1 vs T5 | 1.60 (-1.30, 8.20) | 0.024* | V = 58 | 1.83 (-1.01, 8.11) | 0.010* | V = 81 |
| T2 vs T3 | 0.00 (-2.00, 3.20) | 0.234 | V = 63 | 0.60 (-1.67, 2.17) | 0.035* | V = 86 |
| T2 vs T4 | 0.80 (-4.00, 4.50) | 0.24 | V = 47 | 1.30 (0.03, 2.46) | < 0.001*** | V = 91 |
| T2 vs T5 | 1.00 (-1.00, 4.30) | 0.067 | V = 54 | 1.33 (0.23, 3.45) | < 0.001*** | V = 91 |
| T3 vs T4 | 0.70 (-3.60, 1.90) | 0.365 | V = 44 | 0.44 (0.18, 0.87) | < 0.001*** | V = 91 |
| T3 vs T5 | 0.90 (-2.50, 2.90) | 0.054 | V = 55 | 0.71 (0.21, 1.28) | < 0.001*** | V = 91 |
| T4 vs T5 | 0.40 (-1.80, 4.20) | 0.563 | V = 40 | 0.36 (0.03, 0.99) | < 0.001*** | V = 91 |

**Supplementary Table III:** Within-group comparisons for the time to peak GSR and absolute mean GSR for the nVNS group. In the nVNS group, the time to peak GSR did not change between each successively applied noxious thermal stimulus. The mean GSR measured after each noxious thermal stimulus did not increase from T4 to T5, or from T1 to T2.

| | Time To Peak ($\Delta$) | | | Absolute Mean($\Delta$) | | |
| --- | --- | --- | --- | --- | --- | --- |
|  | Median (Range) | p-value | Test Stat. | Median (Range) | p-value | Test Stat. |
| **T1 vs T2** | 0.35 (-7.30, 17.80) | 0.323 | V = 88 | 0.74 (-2.65, 1.83) | 0.159 | V = 96 |
| **T1 vs T3** | 0.10 (-8.60, 19.80) | 0.528 | V = 81 | 1.01 (-0.91, 3.15) | 0.016* | V = 114 |
| **T1 vs T4** | 0.20 (-8.10, 16.20) | 0.296 | V = 70 | 1.63 (-0.68, 4.77) | 0.003** | V = 110 |
| **T1 vs T5** | 0.10 (-4.70, 14.60) | 0.635 | V = 53 | 1.67 (-0.38, 5.43) | 0.005** | V = 95 |
| **T2 vs T3** | -0.05 (-4.00, 14.70) | 1 | V = 68 | 0.67 (-0.24, 1.97) | < 0.001*** | V = 128 |
| **T2 vs T4** | -0.40 (-8.40, 13.10) | 0.952 | V = 54 | 1.00 (-0.88, 3.36) | 0.001** | V = 113 |
| **T2 vs T5** | -0.20 (-9.50, 14.40) | 0.946 | V = 44 | 1.07 (-0.58, 3.60) | < 0.001*** | V = 101 |
| **T3 vs T4** | 0.25 (-5.90, 12.60) | 0.626 | V = 61 | 0.24 (-0.63, 1.62) | 0.007** | V = 106 |
| **T3 vs T5** | 1.40 (-7.00, 13.90) | 0.34 | V = 60 | 0.52 (-0.34, 2.28) | 0.007** | V = 94 |
| **T4 vs T5** | -0.40 (-4.00, 3.40) | 0.944 | V = 44 | 0.30 (-0.77, 0.66) | 0.268 | V = 71 |

